# Whole Genome Sequencing and Assembly of the Asian Honey Bee *Apis dorsata*

**DOI:** 10.1101/840207

**Authors:** Sara Oppenheim, Xiaolong Cao, Olav Rueppel, Panuwan Chantawannakul, Sasiprapa Krongdang, Patcharin Phokasem, Rob DeSalle, Sara Goodwin, Jinchuan Xing, Jeffrey Rosenfeld

## Abstract

The Asian honey bee (*Apis dorsata)* is distinct from its more widely distributed cousin *A. mellifera* by a few key characteristics. Most prominently, *A. dorsata*, nest in the open by forming a colony clustered around the honeycomb, while *A. mellifera* nest in concealed cavities. Additionally, the worker and reproductive castes are all of the same size in *A. dorsata*. In order to investigate these differences, we performed whole genome sequencing of *A. dorsata* using a hybrid Oxford Nanopore and Illumina approach. The 223MB genome has an N50 of 35kb with the largest scaffold of 302kb. We have found that there are many genes in the *dorsata* genome that are distinct from other hymenoptera and also large amounts of transposable elements, and we suggest some candidate genes for *A. dorsata*’s exceptional level of defensive aggression.

## Introduction

The Asian honey bee (*Apis dorsata)* is an important pollinator and source of honey throughout its natural range in Southeast and East Asia [1–3]. This species is distinct from the European honey bee (*Apis mellifera*) both morphologically and behaviorally. Unlike *A. mellifera*, which is now found throughout the world, *A. dorsata* is found only in Southeast and East Asia [3, 4]. *Apis dorsata* is much larger than *A. mellifera*, and *A. dorsata* workers are almost twice as long as *A. mellifera* workers. In addition, *A. dorsata* is distinguished by a lack of body size variation between castes—in other honey bees, reproductive castes are larger than workers, but in *A. dorsata* all castes are the same size.

The flavor of *A. dorsata* honey differs from that of *A. mellifera*. Biochemical assays of honey from different sources have demonstrated variation in antibacterial activity, antimicrobial activity, protein content, and glucose content [5, 6]. While much of this variation can be explained by the nectar source that bees use to produce the honey, honey characteristics also differ based on honeybee species.

From an anthropocentric point of view, the most striking difference between *A. dorsata* and other honey bees is in their nesting behaviour. *Apis dorsata* do not nest in cavities but instead build exposed nests that hang from tree branches or cliffs. The exposed comb is covered at all times by a blanket of up to 100,000 worker bees. Unlike species that nest in enclosed spaces like tree trunks and rock crevices, *A. dorsata* colonies are unwilling to live in the traditional Langstroth hives used by commercial beekeepers [7]. This distinct lifestyle, together with an exceptional level of defensive aggression [8], has limited the ability of humans to domesticate the species and to commercially produce its honey [7].

To fully understand the genetic basis of important *Apis* traits requires comparative data from a variety of *Apis* species. In this study, we combine short-read and long-read sequencing technologies to generate a high-quality assembly of the *A. dorsata* genome and compare its properties with those of other *Apis* species.

## Materials and Methods

### Sample collection

Multiple *A. dorsata* drones were collected by on the campus of Chiang Mai University (Chiang Mai, Thailand). The drones were collected in the early evening when they visited artificial light sources near the University’s gymnasium. Specimens were sacrificed immediately. One exemplar was kept intact; each of the remaining specimens was cut into 3 sections and placed in RNAlater™ Stabilization Solution (Invitrogen). Tissues were held in RNAlater at 4C for at least 24 hours to allow the solution to saturate the tissue, then shipped to the American Museum of Natural History (AMNH). The exemplar specimen was accessioned into the Ambrose Monell Cryo Collection at AMNH. The remaining specimens were sent to the Woodbury Genome Center at Cold Spring Harbor Laboratories for DNA extraction and sequencing.

### DNA extraction and sequencing

Individual heads and thoraces were chopped into small pieces and placed in 1.7ml centrifuge tubes (~25 mg of tissue per tube). DNA was isolated using the MagAttract HMW DNA kit (Qiagen), following the “Manual Purification of High-Molecular-Weight Genomic DNA from Fresh or Frozen Tissue” protocol. Several tubes were pooled after extraction.

Extracted DNA was sheared to 20kb using a Covaris G-tube following manufacturer’s recommendations. The sheared DNA was prepared for Nanopore sequencing using the Ligation Sequencing 1D kit (SQK-LSK108). Briefly, 1-1.5ug of fragmented DNA was end-repaired with the NEB FFPE repair kit, followed by end repair and A-tailing with the NEB Ultra II end-prep kit. Following an Ampure clean-up step, prepared fragments were ligated to ONT-specific adapters with the NEB blunt/TA master mix kit. After a final Ampure clean-up, the library was loaded on to a MinION R9.0 SpotON flowcell as per manufacturer’s instructions. The flowcell was sequenced with standard parameters for 2 days. The resulting sequence data was processed with the Albacore pipeline (Oxford Nanopore), which performs base calling and other postprocessing steps.

Illumina libraries were prepared with the Kapa hyper prep kit, following manufactures guidelines. Prepared libraries pooled and sequenced on an Illumina NextSeq 500. The sequencing was done with PE150 High Output format.

### Genome Assembly and Annotation

The Nanopore and Illumina reads were assembled together using the SPAdes software [9]. Structural annotation of the assembled genome was performed with MAKER as described in [10]. Briefly, in the first round, transcript sequences from a previous *A. dorsata* study (NCBI BioBroject PRJNA174631) were provided to MAKER as EST evidence. Protein evidence was obtained by collecting protein sequences from six closely related hymenopteran species from NCBI, including *Apis dorsata* (PRJNA174631), *A. mellifera* (PRJNA477511), *A. cerana* (PRJNA324433), *A. florea* (PRJNA45871), *Polistes dominula* (PRJNA307991) and *P. canadensis* PRJNA301748). In addition, all protein sequences in the Swiss-Prot database were included in the analysis [11]. Transcript sequences from the non-*dorsata* species were provided to MAKER as alternative EST evidence. Repetitive elements were identified using RepeatMasker [12].

Coding sequences were predicted in the MAKER pipeline with Exonerate [13] and BLAST [14]. The output of the first MAKER run was used to train the gene predictors SNAP [15] and AUGUSTUS [16]. The trained models were then provided to MAKER and a new set of training models generated from the result. used for another round of running MAKER. Another sets of SNAP and AUGUSTUS models were trained with the new outputs. The assembled contig sequences were used to train GeneMark-ES [17]. The SNAP, AUGUSTUS, and GeneMark-ES models were used in a final MAKER run, and the resulting predicted coding sequences used in all subsequent analyses. The completeness of the genome assembly and MAKER protein models were analysed with BUSCO, which measures completeness in terms of evolutionarily informed expectations of gene content [18]. The BUSCO Insecta dataset (1,658 single-copy conserved genes) was used as a reference.

### Functional annotation

Annotation of the predicted genes was undertaken with a variety of tools. First, we conducted BLASTp searches against all Hymenoptera entries in the NCBI nr database [19]. For sequences that had no hit, we did a second BLASTp search against all sequences in the NCBI nr database. We used InterProScan 5 [20] to classify genes into protein families and identify functional domains. InterProScan runs many different tools to compare query sequences to the InterPro [21] protein signature databases. The BLASTp and InterProScan results were imported into Blast2GO [22], where we mapped them to Gene Ontology (GO) terms [23].

We also compared the content of the assembled *A. dorsata* genome with that of *A. mellifera* (which is the most complete *Apis* genome available).

### Ortholog identification

We used OrthoPipe, a stand-alone pipeline version of OrthoDB 2.3.1 [24], to classify the protein-coding genes from *A. dorsata* and three other bee species with similarly complete genome data available (*A. mellifera*, *A. cerana*, *A. florea*) into orthologous clusters. To evaluate the resulting clusters in a phylogenetic context, we used the reference topology presented by [25] to identify clusters at each internal node that contained all taxa subtended by that node.

### *Identification of species-specific* A. dorsata *genes*

Within the context of these analyses, species-specific genes were narrowly defined as sequences that lacked orthologs in any other species in this study (i.e., occurred only at terminal nodes). A more stringent definition requires a lack of orthologs in any published data set [26], and we used BLASTp searches against NCBI’s nr database to test whether our narrowly defined species-specific genes were legitimate orphans.

### Functional enrichment tests

Because raw frequency-based comparisons are difficult to interpret in evolutionary terms, we used enrichment tests to examine whether any gene functions were over-represented in the genome of *A. dorsata* as compared to *A. mellifera*. We conducted Fisher’s exact tests (False Discovery Rate adjusted p-value < 0.01) to identify GO terms or InterPro signatures enriched in a test set of annotated genes relative to a reference set. Since GO annotations, which are assigned transitively via BLAST hits, can sometimes lead to incorrect functional annotation [27], we used InterPro signatures as the primary source of functional annotation and comparison.

## Results and Discussion

### Genome assembly

Our assembly results were consistent with previous Apis genome sequencing projects (Figure 1): the total assembly size was 224 Mb (versus 225 to 230 Mb in other apid bees [28–30]), with a scaffold N50 of 35 kb (largest scaffold: 302 kb). BUSCO analysis showed that the assembly was 98.9% complete. We identified a wide variety of repetitive elements (Supplemental Table 1), and these comprised ~7% of the genome. Following the Maker pipeline described above, we identified 13,517 protein coding genes (versus 12,145 to 12,940 genes in other apids [28–30]).

**Figure 1.**
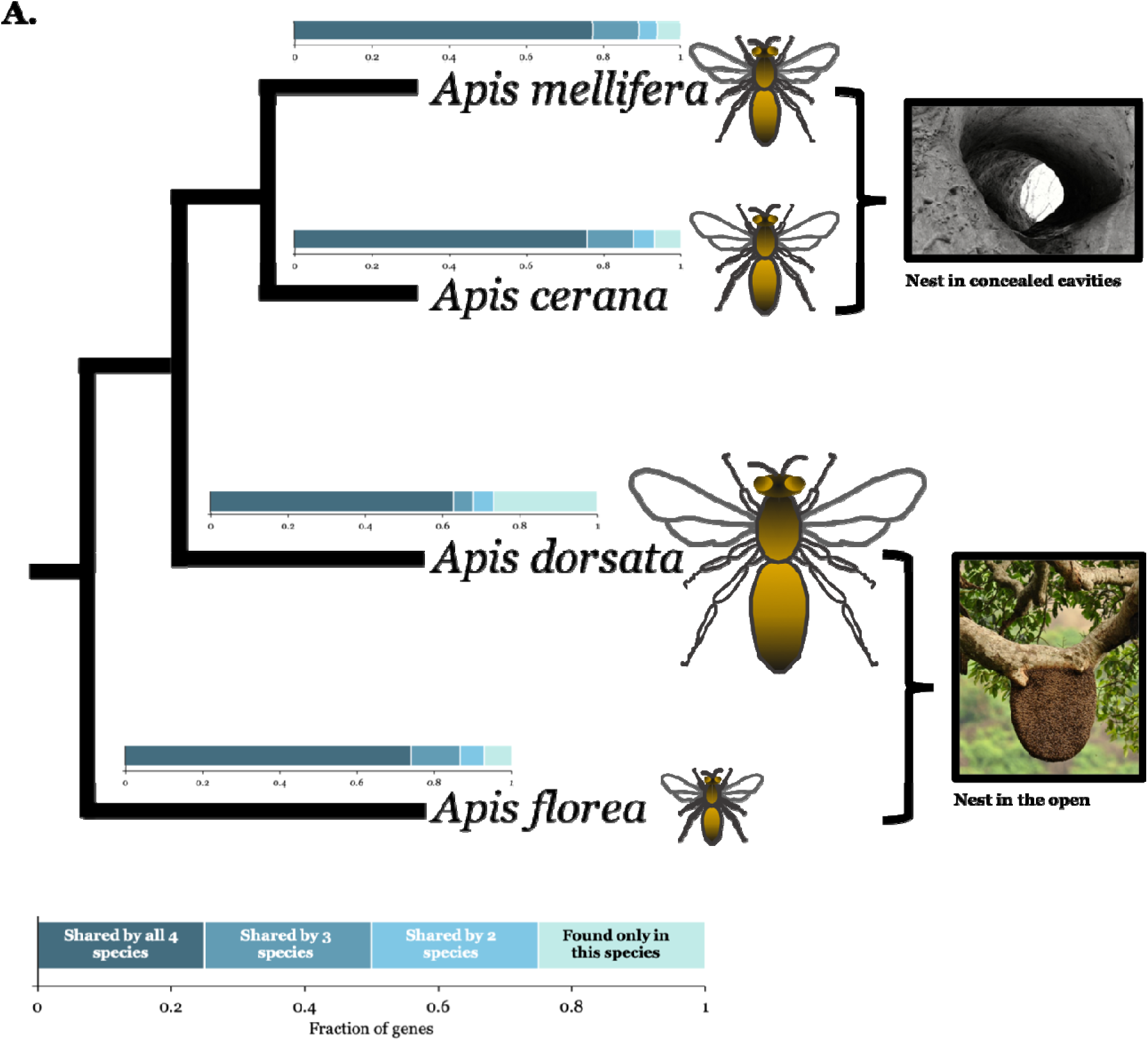
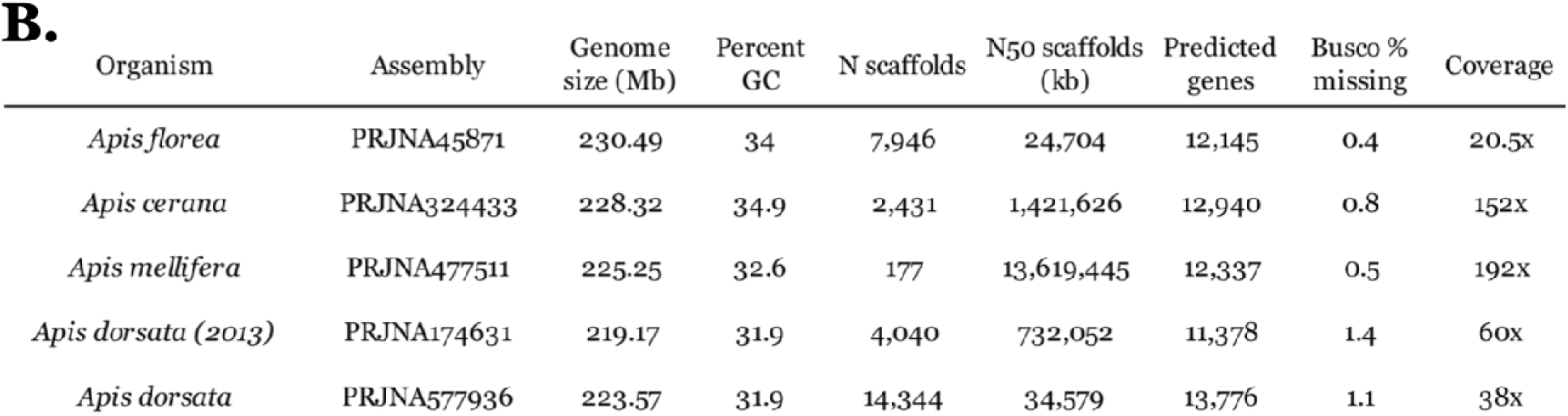
**A)** Phylogenetic relationship between the four *Apis* species used for ortholog identification, relative sizes of drones are shown. Inset bar charts indicate the fraction of genes from each species that were orthologous at different levels: universal orthologs (present in all four species), species-specific genes (those found only in one species), and others. **B)** Assembly statistics for *Apis* species.

### Functional annotation

More than 80% (11,264) of the predicted *A. dorsata* genes had hits to other Hymenopteran sequences in the nr database. The remaining 2,253 genes had no hits to other hymenoptera (or only had hits to *A. dorsata* sequences). When we blasted these genes against the entire nr database, 182 of them had hits but 2,071 still had no hit. The results of InterProScan analyses were similar: 1,956 genes had no InterPro signature. The overlap between these annotations left us with 1,251 genes that lacked functional annotation from either blast or InterProScan.

### Ortholog identification

We identified 3,451 single copy universal ortholog clusters that contained one gene from each of the four species included in ortholog analysis. Consistent with the idea that single copy universal orthologs represent highly conserved genes, every *A. dorsata* single copy universal ortholog had a hit to other Hymenoptera in the nr database. Seventy-three percent of the *A. dorsata* genes (9,900) had orthologs in at least one other species, versus 93% of the *A. mellifera* genes. In the specific comparison between *A. mellifera* and *A. dorsata*, 22% (3,034) of the *A. dorsata* genes had no ortholog in *A. mellifera*, and 2,177 of these had no ortholog in any of the included *Apis* species and no hit to any other hymenopteran in our blast searches (Figure 1).

### *Species-specific* A. dorsata *genes*

Some 3,617 *A. dorsata* genes did not have orthologs in any of the included *Apis* species. We used blast searches against all hymenopteran sequences in the nr database to determine whether these were species-specific genes. We found that 1,424 of these genes had a blast hit to another hymenopteran species, and an additional 75 had a hit to a non-hymenopteran species. This left 2,118 genes that appear to be either pseudogenes or species-specific *A. dorsata* genes because they had no hit to any sequence in the nr database.

### Divergence in 5-hydroxytryptamine receptor 2A (5-HT2a)

We were particularly interested in identifying genes that might contribute to *A. dorsata*’s behavioural divergence from other *Apis* species. We focused on receptors for the neurotransmitter serotonin (5-hydroxytryptamine, 5-HT) because of serotonin’s role as a neuromodulator [31]. The effects of serotonin depend on receptor-specific binding, and in *Drosophila* and other insects 5-HT receptors have been associated with variation in locomotion, feeding behavior, learning and memory, and aggression [32, 33].

We identified four 5-HT2A receptors in *A. dorsata* (Figure 2A-B). While three of these had orthologs in other *Apis* species, one was sufficiently divergent that it did not cluster with any other *Apis* genes. Because serotonin has a stimulatory effect on aggression in some insects [34], this gene is an attractive candidate for future studies on the genetic basis of aggression in *A. dorsata*.

**Figure 2.**
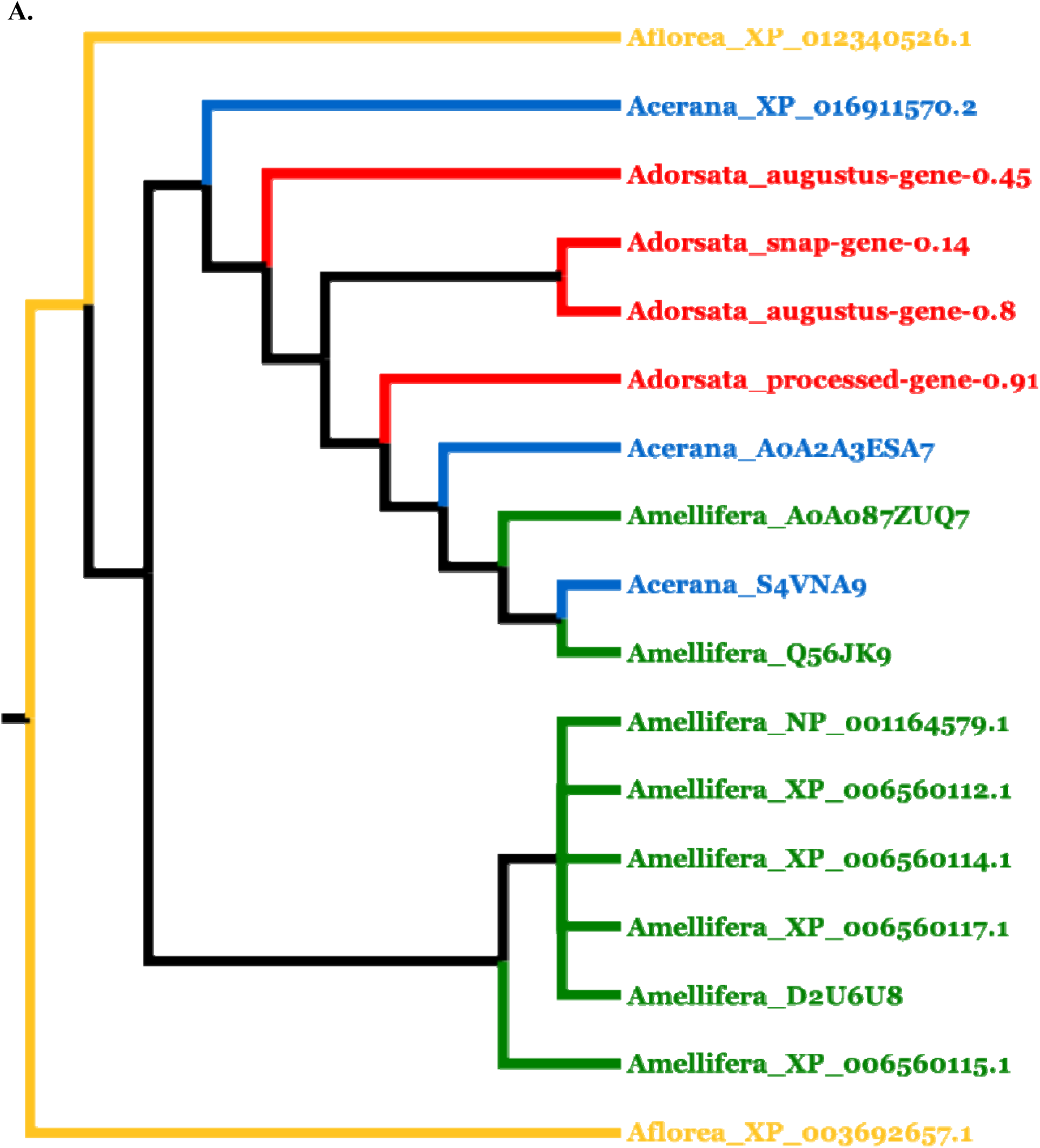
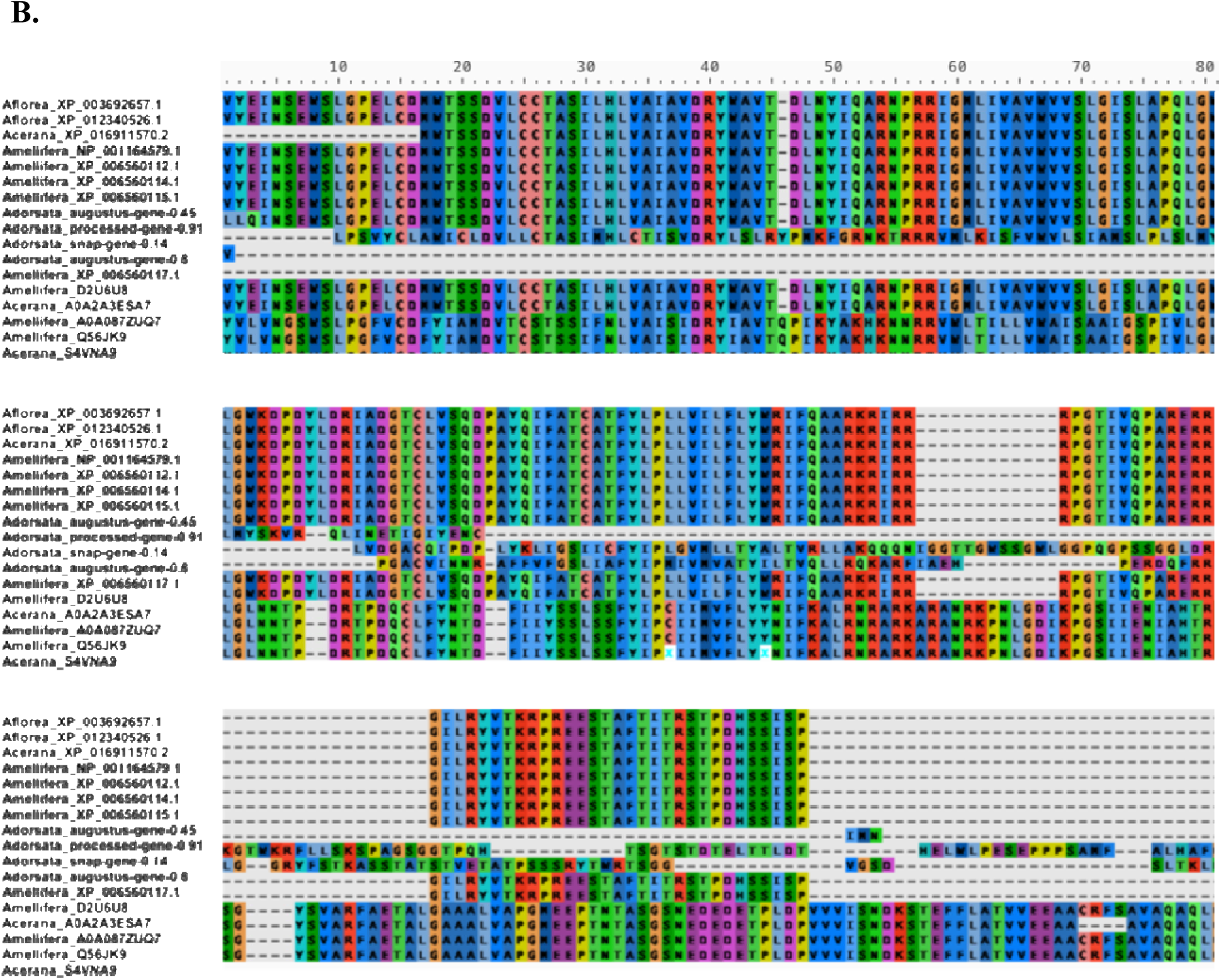
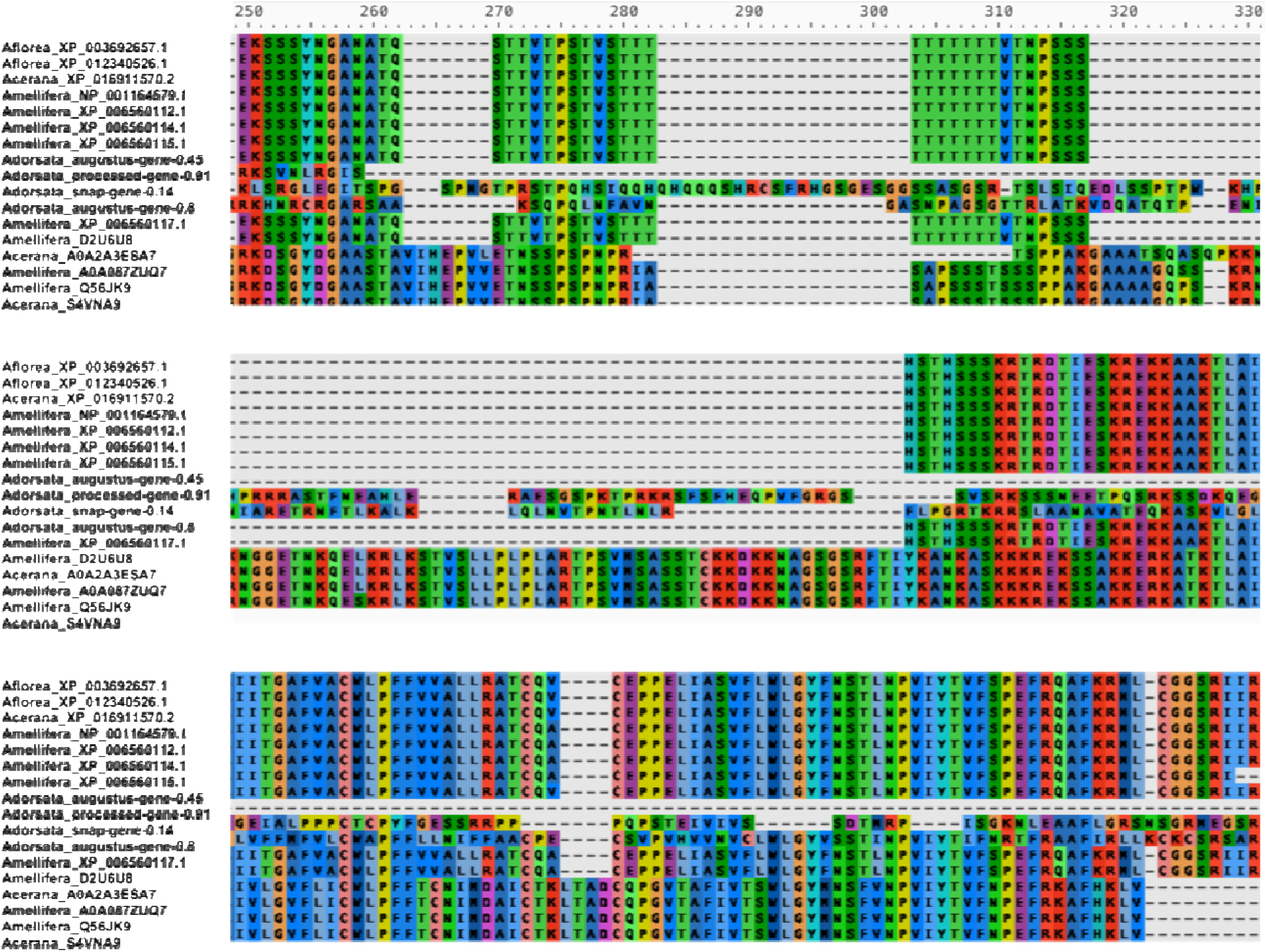
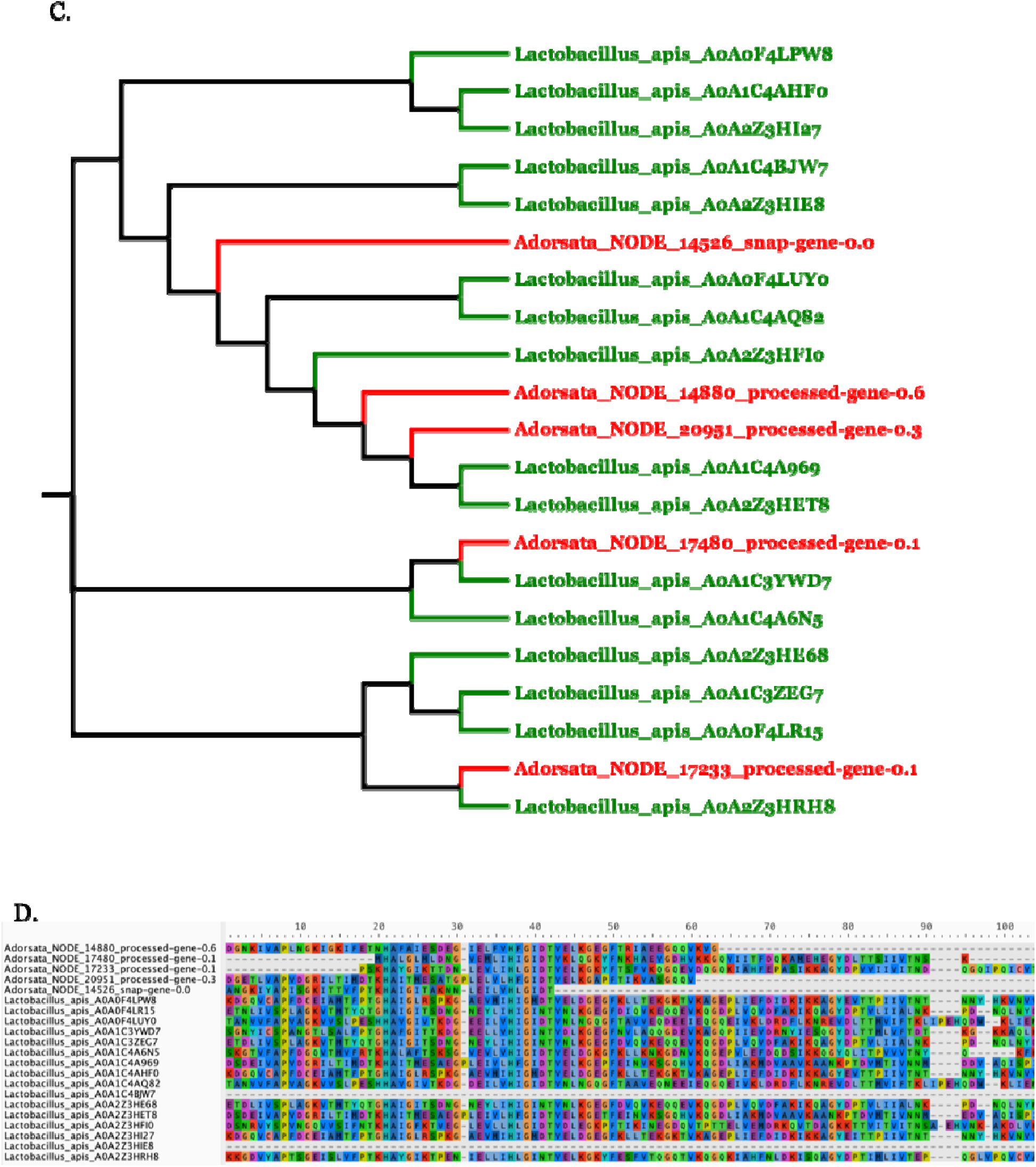
**A)** Cladogram and **B**) sequence alignment of putative 5-hydroxytryptamine (serotonin) receptor 2A (5-HT2A) genes from *A. dorsata* and other *Apis* species. **C)** Cladogram and **D)** Sequence alignment of *A. dorsata* species-specific genes with InterPro signature IPR001127 (Lactobacillus PTS carbohydrate transport). *Lactobacillus apis* genes are from the gut of *A. mellifera*.

### Functional enrichment

#### Single copy universal orthologs versus all other genes

These highly conserved genes were enriched for functions related to development, including a growth factor responsible for cell proliferation and differentiation (IPR001111), and juvenile hormone binding proteins (JHBP; IPR010562), which regulates embryogenesis, larval development, and reproductive maturation in the adult forms (Supplemental Table 2).

#### *Species-specific* A. mellifera *genes versus all other genes*

The *A. mellifera* genes that lacked *A. dorsata* homologs were enriched for signal transduction functions. Several enriched signatures related to small GTPases, which are critical components in cellular signal transduction pathways,

#### *Species-specific* A. dorsata *genes versus all other genes*

The InterPro terms that were enriched in the *A. dorsata* genes that lacked *A. mellifera* homologs fell into two main functional categories. First, there were 28 genes with Ty1/copia-like retrotransposon signatures. The “domestication” of transposable elements can lead to the emergence of species-specific genes by providing a source of biochemically active elements such as transcription factor-binding sites, and by generating genomic rearrangements [35–38], and the over-representation of transposable element-related signatures in the genome of *A. dorsata*, suggests that the evolution of “new” genes may have played a role in the behavioral diversification of *A. dorsata* from other honeybees.

Second, there were 23 sequences that appear to be part of *A. dorsata*’s microbiome assemblage. These all had high-scoring hits to bacterial species in the nr database, and when aligned with bacterial sequences known to occur in Hymenopteran microbiomes showed high levels of similarity. The enriched InterPro signature IPR001127, a domain that is associated with the carbohydrate transport system in *Lactobacillus* bacteria, was found in six of the species-specific *A. dorsata* genes. We compared the *A. dorsata* sequences to *Lactobacillus apis* genes from the microbiome of *A. mellifera* (Figure 2C-D).

In honeybees, the microbiome of worker bees is of especial importance. All worker gut microbiomes studied to date contain abundant carbohydrate-processing genes, but the exact components of the microbiome community show variation between honeybee strains and species [39]. This variation suggests that some of the dietary differences between honeybees in different localities (in their use of pollen/nectar from different plant species) might be controlled by microbiome components [40–42]. Previous research has shown that *Lactobacillus* species, which were the most common top hits for our *A. dorsata*-only sequences, play a major role in carbohydrate digestion [43, 44]. Thus, the specific microbiome components present in different colonies or species might determine their nutritional ecology and/or their ability to cope with dietary toxins and could explain the geographic distribution of *A. dorsata* as a case of coevolution between bees, bacteria, and flowering plants.

The microbiome of *A. dorsata* may also be involved in the distinctive flavour of the honey they produce. *Lactobacillus* species are thought to prevent the fermentation of stored honey by retarding the growth of fermenting yeasts [45]. Variation in the activity of *Lactobacillus* strains in honeybee microbiomes can affect the flavour of honey since flavour is partially dependent on lactic acid bacterial metabolites [46]. Thus, the “terroir” of different honeys [47] may result as much from the honeybee microbiome [48] as from the particular locale where the honeybees forage.

## Acknowledgements

This research work was partially supported by Chiang Mai University. This research used resources from the Rutgers Discovery Informatics Institute, which are supported by Rutgers and the State of New Jersey [49]. Some of this work was conducted on the BIOMIX compute cluster, made possible through funding from Delaware INBRE (NIGMS P20GM103446), the State of Delaware, and the Delaware Biotechnology Institute. Dr. Jeffrey Rosenfeld is supported by Cancer Center Support grant (2P30-CA072720-20) and the Systems Biology program at the Cancer Institute of New Jersey. We would like to thank Corey Smith at the American Museum of Natural History for Assistance in navigating the government paperwork for the import of the specimens.

## Data availability

Genome assembly and raw sequencing reads are available as part of NCBI PRJNA577936.

