## Supplementary Tables for "Whole Genome Sequencing and Assembly of the Asian Honey Bee *Apis dorsata*"

**Supplemental Table 1.** RepeatMasker results.

| Type | N elements | Length (bp) | Percent of sequence |
|---|---|---|---|
| **Retroelements** | 5,004 | 360,562 | 0.16% |
| **SINEs** | 20 | 1,919 | 0.00% |
| Penelope | 658 | 46,476 | 0.02% |
| **LINEs** | 2,115 | 154,766 | 0.07% |
| L2/CR1/Rex | 73 | 3,824 | 0.00% |
| R1/LOA/Jockey | 28 | 2,514 | 0.00% |
| R2/R4/NeSL | 13 | 1,250 | 0.00% |
| RTE/Bov-B | 208 | 19,795 | 0.01% |
| L1/CIN4 | 737 | 50,727 | 0.02% |
| **LTR elements** | 2,869 | 203,877 | 0.09% |
| BEL/Pao | 265 | 24,018 | 0.01% |
| Ty1/Copia | 31 | 7,678 | 0.00% |
| Gypsy/DIRS1 | 838 | 62,256 | 0.03% |
| Retroviral | 1,218 | 70,865 | 0.03% |
| **DNA transposons** | 11,155 | 748,965 | 0.33% |
| hobo-Activator | 3,206 | 204,429 | 0.09% |
| Tc1-IS630-Pogo | 632 | 102,005 | 0.05% |
| PiggyBac | 90 | 4,603 | 0.00% |
| Tourist/Harbinger | 794 | 43,897 | 0.02% |
| Other (Mirage, P-element, Transib) | 139 | 6,220 | 0.00% |
| **Rolling-circles** | 425 | 27,897 | 0.01% |
| **Unclassified** | 1,387 | 156,892 | 0.07% |
| **Total interspersed repeats** | . | 1,266,419 | 0.57% |
| **Small RNA** | 277 | 32,089 | 0.01% |
| **Satellites** | 660 | 68,592 | 0.03% |
| **Simple repeats** | 247,811 | 11,358,037 | 5.08% |
| **Low complexity** | 50,265 | 2,718,971 | 1.22% |
| **Total bases masked** | **.** | **15,396,268** | **6.89%** |

**Supplemental Table 2.** Enriched InterPro signatures.

| Test set | Reference set | Enriched InterPro signature | InterPro name | Functional importance | FDR | p-value | N test set | N ref set |
|---|---|---|---|---|---|---|---|---|
| *A. dorsata* -only genes | All *A. dorsata* and *A. mellifera* genes | IPR010992 | Integration host factor (IHF)-like DNA-binding domain | microbiome | 0.01874 | 8.92E-06 | 5 | 0 |
| *A. dorsata* -only genes | | IPR001127 | bacterial phosphoenolpyruvate: sugar phosphotransferase system (PTS) | microbiome | 0.01874 | 5.59E-06 | 6 | 1 |
| *A. dorsata* -only genes | | IPR001584 | Integrase, catalytic core | transposable element | 0.002401 | 3.12E-07 | 8 | 2 |
| *A. dorsata* -only genes | | IPR029526 | PiggyBac transposable element-derived protein | transposable element | 0.01874 | 6.80E-06 | 8 | 5 |
| *A. dorsata* -only genes | | IPR039537 | Retrotransposon Ty1/copia-like | transposable element | 0.004027 | 8.71E-07 | 6 | 0 |
| Single copy universal genes | | IPR000618 | Insect cuticle protein | growth and development | 3.73E-04 | 6.61E-07 | 39 | 58 |
| Single copy universal genes | | IPR001111 | Transforming growth factor beta 1 | growth and development | 8.23E-05 | 1.10E-07 | 12 | 2 |
| Single copy universal genes | | IPR001839 | Transforming growth factor-beta, C-terminal | growth and development | 8.23E-05 | 1.04E-07 | 13 | 3 |
| Single copy universal genes | | IPR010562 | Haemolymph juvenile hormone binding | growth and development | 8.46E-05 | 1.17E-07 | 20 | 13 |
| Single copy universal genes | | IPR001623 | DnaJ domain | molecular chaperone | 2.22E-05 | 1.92E-08 | 35 | 39 |
| Single copy universal genes | | IPR002423 | Chaperonin TCP-1, | molecular chaperone | 9.24E-05 | 1.36E-07 | 16 | 7 |
| Single copy universal genes | | IPR011032 | GroES-like superfamily | molecular chaperone | 1.95E-05 | 1.10E-08 | 20 | 10 |
| Single copy universal genes | | IPR006689 | Small GTPase superfamily, ARF/SAR type | signal transduction | 0.009376 | 4.79E-05 | 19 | 22 |
| *A. mellifera*-only genes | | IPR000159 | Ras-associating (RA) domain | signal transduction | 8.44E-08 | 3.62E-10 | 21 | 81 |
| *A. mellifera*-only genes | | IPR001478 | PDZ | signal transduction | 2.16E-07 | 9.71E-10 | 46 | 386 |
| *A. mellifera*-only genes | | IPR000651 | Ras-like guanine nucleotide exchange factor, N-terminal | signal transduction | 6.13E-05 | 3.32E-07 | 11 | 30 |
| *A. mellifera*-only genes | | IPR000014 | PAS | signal transduction | 3.34E-04 | 2.22E-06 | 8 | 16 |
| *A. mellifera*-only genes | | IPR001895 | Ras guanine-nucleotide exchange factors catalytic domain | signal transduction | 3.36E-04 | 2.30E-06 | 11 | 38 |
| *A. mellifera*-only genes | | IPR000331 | Rap GTPase activating protein domain | signal transduction | 9.92E-04 | 7.30E-06 | 10 | 35 |
| *A. mellifera*-only genes | | IPR002073 | 3'5'-cyclic nucleotide phosphodiesterase, catalytic domain | signal transduction | 0.001984 | 1.63E-05 | 10 | 39 |
